# The interplay between information flux and temporal dynamics in infraslow frequencies

**DOI:** 10.1101/2020.06.11.106476

**Authors:** Mehrshad Golesorkhi, Shankar Tumati, Javier Gomez-Pilar, Emmanuel. A. Stamatakis, Georg. Northoff

## Abstract

Unlike the brain’s faster frequencies, the exact role of its more powerful infraslow frequencies (ISF, 0.01 – 0.1Hz) in information processing remains poorly understood. Do and how ISF process information? We investigate information processing and related temporal dynamics of ISF in resting and task state fMRI. To quantify information, we apply the Lempel-Ziv complexity (LZC), a measure of signal compression indexing information. The LZC is combined with direct measurement of the dynamics of ISF themselves, namely their power spectral density by median frequency (MF). We demonstrate the following: (I) topographical differences in resting state between higher- and lower-order networks, showing statistically lower LZC in the former; (II) task-related changes in LZC; (III) modulation of LZC associated with MF changes, with low and high MF resting-state values correlated with different degrees of LZC change. In sum, we provide evidence that ISF carry and process information as mediated through their temporal dynamics.

## Introduction

The brain exhibits fluctuations in its neural activity which includes different frequency ranges. We know very well that the brain’s faster frequencies in the range of 1 to 240Hz process information and thereby shape behavior and cognition as measured in EEG/MEG^1–5^. Unlike its faster frequencies, the function of the brain’s very slow waves, its infraslow frequencies (ISF, 0.01 - 0.1Hz) is less understood ^1,6^. ISF are often conceived as noise stemming from vascular sources^7,8^ rather than carrying information. However, recent evidence hints upon a neurophysiological basis of ISF ^9–20^, which suggests their potential involvement in processing information. Do ISF, like the brain’s faster frequencies, process information during resting and task states? If so, how do ISF process information, i.e., through which pattern? The goal of our paper is to address these two questions. For that purpose, we probe “whether” ISF carry information by investigating them in fMRI with a well-established measure from the Information Theory field, i.e., Lempel-Ziv Complexity (LZC) (see below). To test “how” ISF process information, we combine LZC with a measurement of the dynamics of ISF, namely their power spectral density by median frequency (MF) (see below).

ISF are predominantly measured with functional magnetic resonance imaging (fMRI) that is based on the blood oxygen level-dependent (BOLD) signal ^21,22^. Some conceive the BOLD signal to reflect vascular low-pass filtering of higher frequency neural activity – in that case, the observed ISF would be at best vascular noise without a distinct neurophysiological basis. That is contradicted by recent results which demonstrate a specific neurophysiological basis of ISF ^9–20^. The neurophysiological basis of ISF is further supported by their occurrence in particular cortical layers ^6,10^ as well as by various electrophysiological features like phase-specific distributions and coherence ^9,10,13,14^, local field potentials (LFP) ^11,12,15,17–21^, and power spectrum ^9^. Moreover, task-related changes in ISF have been reported for instance during attention ^23,24^ and conscious perception ^25^. Together, such distinct neurophysiological basis of ISF suggests that they are not mere noise but carry and process information. However, “whether” the ISF do, in fact, carry information remains yet to be demonstrated.

Various fMRI studies demonstrated topographical organization of ISF with lower-order networks, i.e., sensory networks, and higher-order networks, i.e., default-mode network, executive network, and others, standing at opposite ends of the spatial ^26–29^. Interestingly, such a spatial hierarchy converges with a corresponding temporal hierarchy along the continuum of fast-slow timescales ^9,26–28,30–34^. These findings are mostly based on functional connectivity (FC) ^26,29^ and intrinsic neural timescales as measured by autocorrelation window (ACW) in fMRI resting state ^33^. They thus describe the spatial and temporal topographical organization of the brain’s ISF.

This leaves open whether the ISF themselves including their spatial and temporal organization carry and process information. For that, one needs to operationalize information which, for instance, is provided by Lempel-Ziv Complexity (LZC) ^35–37^: the LZC measures the number of distinct patterns in a binary sequence, namely the degree to which a signal can be compressed ^38,39^. It reflects the amount of information (number of bits) required to reconstruct a signal ^38^ (see the validity of LZC in fMRI ^40–43^ as well as in other imaging modalities including MEG, EEG, and spike train analysis ^44–52^). Accordingly, applying the LZC allows probing “whether” the ISF carry information other than simply noise. Ideally, one wants to combine the LZC with a measure of the ISF themselves like their power spectral density, indexed by its median frequency (MF). In more general terms, MF measures the speed (or velocity) of the signal in a specific frequency range (e.g. the ISF) ^53–57^ by characterizing the distribution of its power spectrum. This allows probing “how” the ISF process information (see Fig. 1 for general overview).

**Fig. 1.**
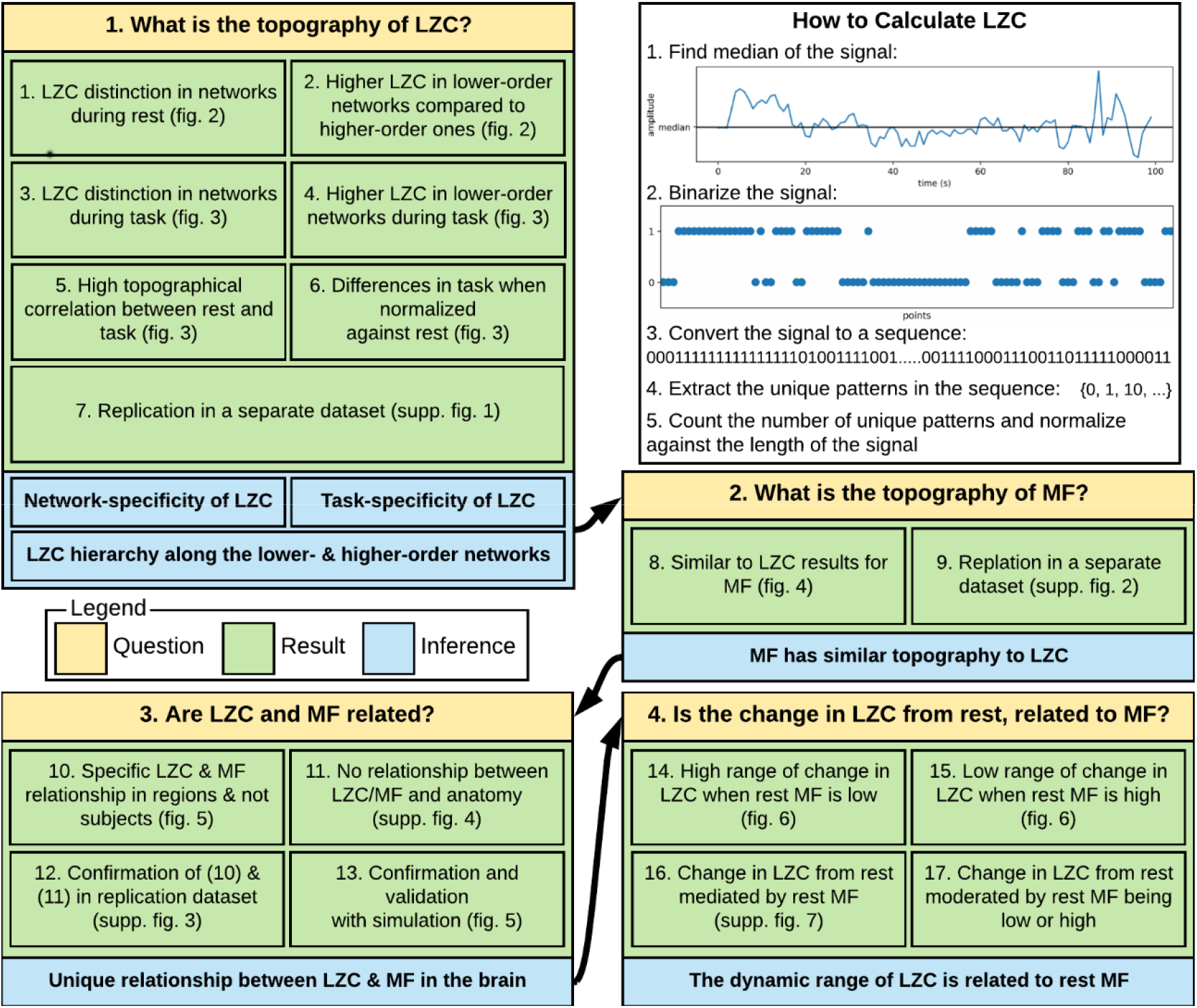
Schema of the paper. Each box is one of the questions being investigated in this work. Each box is divided into three parts: a question (yellow), several analyzes (green), and one or more inferences (blue). The white box contains an overall view of how Lempel-Ziv Complexity (LZC) is calculated.

### Specific aims and hypotheses

The first specific aim is to investigate the LZC and MF of ISF in the resting state. This addresses our first key question namely “whether” ISF carry information. Our first hypothesis is based on the brain’s topography namely, its spatial and temporal hierarchies: if the ISF carry information, we expect the LZC to exhibit different values in the resting state of the lower- and higher-order regions/networks. As most likely related to their continuous exposure to high loads of fast external information input ^26,28,58–60^, lower-order regions like sensory networks and multimodal sensory regions may show higher LZC and faster MF. While higher-order regions/networks, i.e., default-mode network (DMN), frontoparietal network (FPN), language network, and dorsal attention network do not receive such direct information input, but a “filtered” version of it ^26,58,59^. For that reason, we expect lower LZC and lower MF in the high-order networks.

The second specific aim is to investigate changes in LZC and MF of ISF during different task states including a movie and a retinotopy task (as provided by 7T HCP). This serves the purpose to substantiate the first key question, namely that ISF carry information. Since the information load changes from resting to task state, we expect rest-task differences in LZC along the lines of lower- and higher-order networks as both are known to process different loads and kinds of information during task states ^40,42,46,47,61^.

The third specific aim consists of linking LZC to the measurement of the ISF themselves, that is, their power spectral density as indexed by MF. This addresses our second key question, namely “how” the ISF process information. Given that we assume LZC to follow the spatial hierarchy of lower- and higher-order networks and MF to index a corresponding temporal hierarchy, we hypothesize close relation of rest and rest-task modulation of LZC to MF during rest and task states. To establish the LZC-MF relationship, we calculate different correlation and regression models. Furthermore, we conduct various simulation models applying different kinds (pink, white, etc.) and levels of noise to demonstrate that the LZC-MF relationship is specifically related to the brain’s topography rather than noise. This serves the purpose to connect LZC, which measures “whether” ISF carries information, to MF that allows probing “how” the ISF process information: through power spectral density indexing their ‘velocity’ or ‘speed’.

## Results

The main aim of this article is to address our two key questions namely to investigate “whether” and “how” infra-slow frequencies (ISF) carry and process information during resting and different task states (see Fig. 1 for an overview of our guiding questions and their results). We addressed these two key questions by splitting them into a sequence of four different questions as explicated in our overview in Figure 1. Using the 7T of HCP, fMRI signals of 146 subjects during resting and two different task states were parcellated into 717 brain regions defined in the template provided by Pappas and colleagues ^62^. These regions were divided into 12 networks of Visual1, Visual2, Auditory, Somatomotor, Posterior Multimodal, Ventral Multimodal, Orbito Affective, Dorsal Attention, Language, Cingulo Opercular, Frontoparietal and Default. They were also divided into two categories of lower- and higher-order regions. Regions in Visual1, Visual2, Auditory, Somatomotor, Posterior Multimodal, Ventral Multimodal and Orbito Affective networks were put into the lower-order and the rest into the higher-order categories.

To operationalize and measure information, we used the Lempel-Ziv complexity (LZC) which is algorithmically defined as the number of different patterns in a binary sequence indexing the complexity of the BOLD signal (the white box of Fig. 1 shows how LZC is calculated). LZC is formally defined as an index of how much a signal can be compressed, and in other words, it measures how “diverse” are the patterns that are present in a signal, i.e. higher LZC is associated with lower compressibility and more information ^39,63^. The amplitude of regional signals was individually binarized using its median as a robust threshold ^39,64^ and then fed into the LZC algorithm yielding a single LZC value per region of a subject during a specific condition (resting or task state).

Preprocessed 7T fMRI data (TR=1) were downloaded from the Human Connectome Project (HCP) WU-Minn HCP 1200 subjects data release, and then high-pass filtered at 0.1 Hz (more detail about the data release is provided in the method section). The data includes resting state (REST, 16 minutes), and two task states of movie-watching (MOVIE, 15 minutes) and retinotopy (RET, 5 minutes).

### The topography of LZC during the resting state

Our first question is to investigate the spatial distribution of LZC in the resting state (see below for simulation on the validity of LZC measurement in fMRI). After averaging over subjects, the regional distribution of LZC values (Fig. 2A) suggested a specific spatial pattern of LZC. To investigate that, we used the 12 predefined networks (Fig. 2B) and divided the LZC values into them (Fig. 2C). This revealed different LZC patterns among the networks. Performing a one-way analysis of variance (ANOVA) over the 12 networks across regions showed significant () differences among them ().

**Figure 2.**
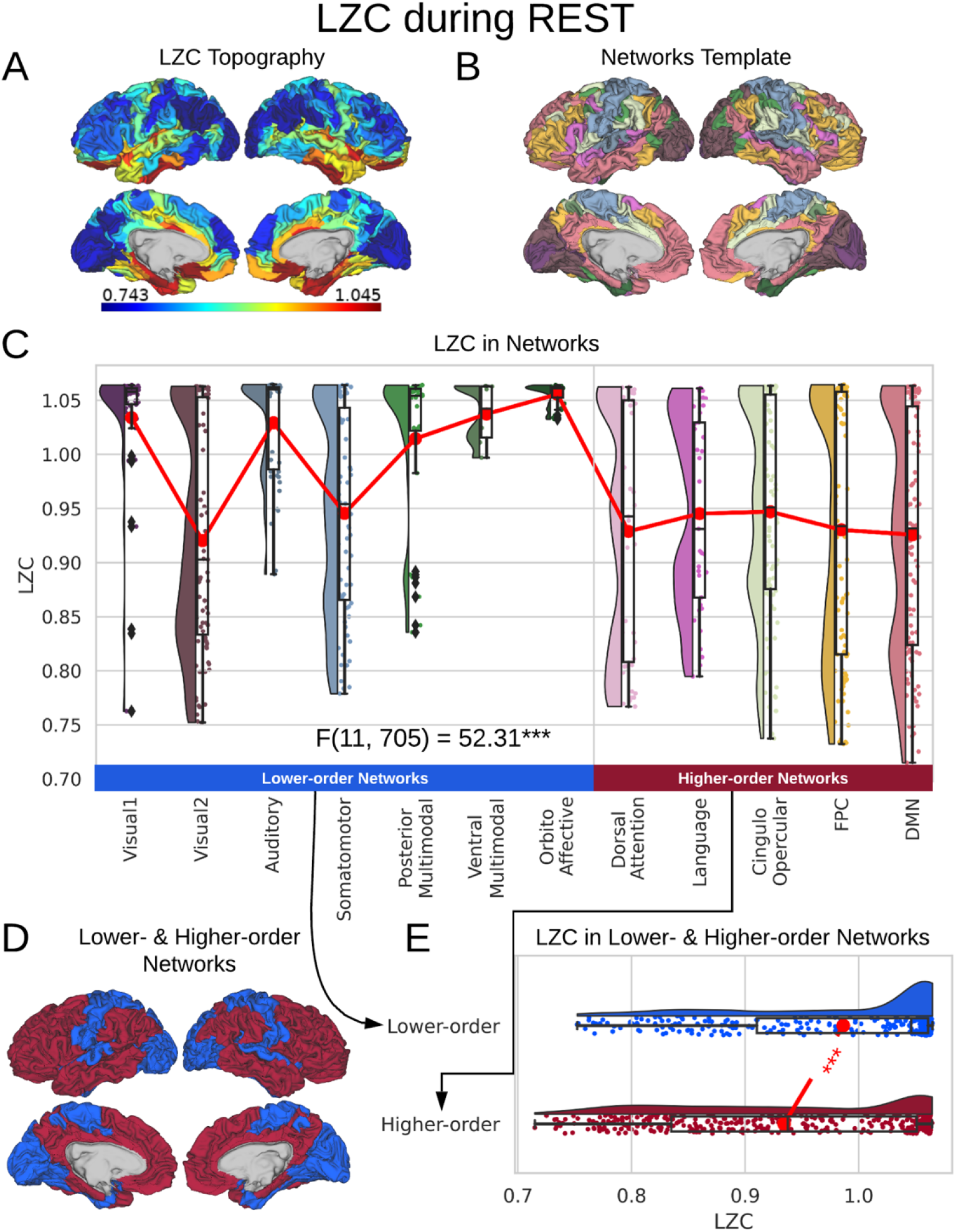
LZC during the resting state. Rainclouds represent regions. **(A)** Spatial distribution of LZC during REST condition. LZC is calculated for a region after binarizing its BOLD signal’s amplitude using the median as a threshold. **(B)** Brain map of the 12 networks defined in ref ^62^. Colors represent different networks. **(C)** Distribution of LZC values in different networks. Colors are the same as in B. Higher-order networks show lower LZC compared to lower-order ones. One-way analysis of variance (ANOVA) showed a significant (p < 0.001) difference among all networks (*F*(11, 705) = 52.31, *η*^2^ = 0.44). **(D)** Brain map representing lower- and higher-order categories. **(E)** LZC during resting state along the lower- and higher-order categories. The student’s t-test shows lower-order regions have significantly higher LZC compared to higher-order ones (*t* = 6.80, *d* = 0.51, *p* < 0.001). Stars represent the significance level (*** ≡ *α* = 0.001)

Significant differences in LZC values among networks paved the way to further investigate the topography of LZC along the lines of lower- and higher-order categories (Fig. 2D). Again, the resting state LZC was averaged over subjects and then compared between lower- and higher-order regions (Fig. 2E). Student’s t-test (with the Cohen’s d for the effect size) showed a significant difference between the two types of networks () with the lower-order having higher LZC compared to the higher-order category. Together, these results show clear topographical differences in the spatial distribution of LZC (These results were also replicated in our 3T replication dataset in Supp. Fig. 1). Given that lower- and higher-order networks are known to process different kinds and degrees of information ^65^, our result of LZC following these topographical differences supports the assumption that ISF carry information. This provides a positive answer to our first key question “whether” ISF carry information.

### The topography of LZC during task states

Spatial distribution of LZC was investigated during the two tasks of movie-watching (MOVIE) and retinotopy (RET) in three different ways: analysis of absolute values, spatial correlations, and analysis of percentage changes, i.e., rest-task difference. The two tasks have different complexity and temporal structure; one containing rich stimuli of movie clips viewed in long intervals and one containing very simple retinotopic stimuli viewed in short intervals. The different input structure of the two tasks suggests that they should impact the information processing in lower- and higher-order networks in different ways; showing that would further substantiate the assumption that ISF carry information.

Similar to our resting state analysis, LZC values were calculated for lower- and higher-order networks (Fig. 3A) and their difference was statistically tested across regions. First, a two-way ANOVA with network order (2 levels: lower vs. higher) and task condition (2 levels: MOVIE vs. RET) as factors was conducted on the LZC values over the regions. The model showed a significant (*p* < 0.001) effect of network order on LZC (*F*(1, 1430) = 27.75, *η*^2^ = 0.02). Further analysis using student’s t-test for each task condition, confirmed that lower-order networks have significantly higher LZC compared to higher-order ones in both MOVIE (t = 4.36, *d* = 0.32, *p* < 0.01) and RET (*t* = 3.22, d = 0.24, *p* < 0.01).

**Fig. 3.**
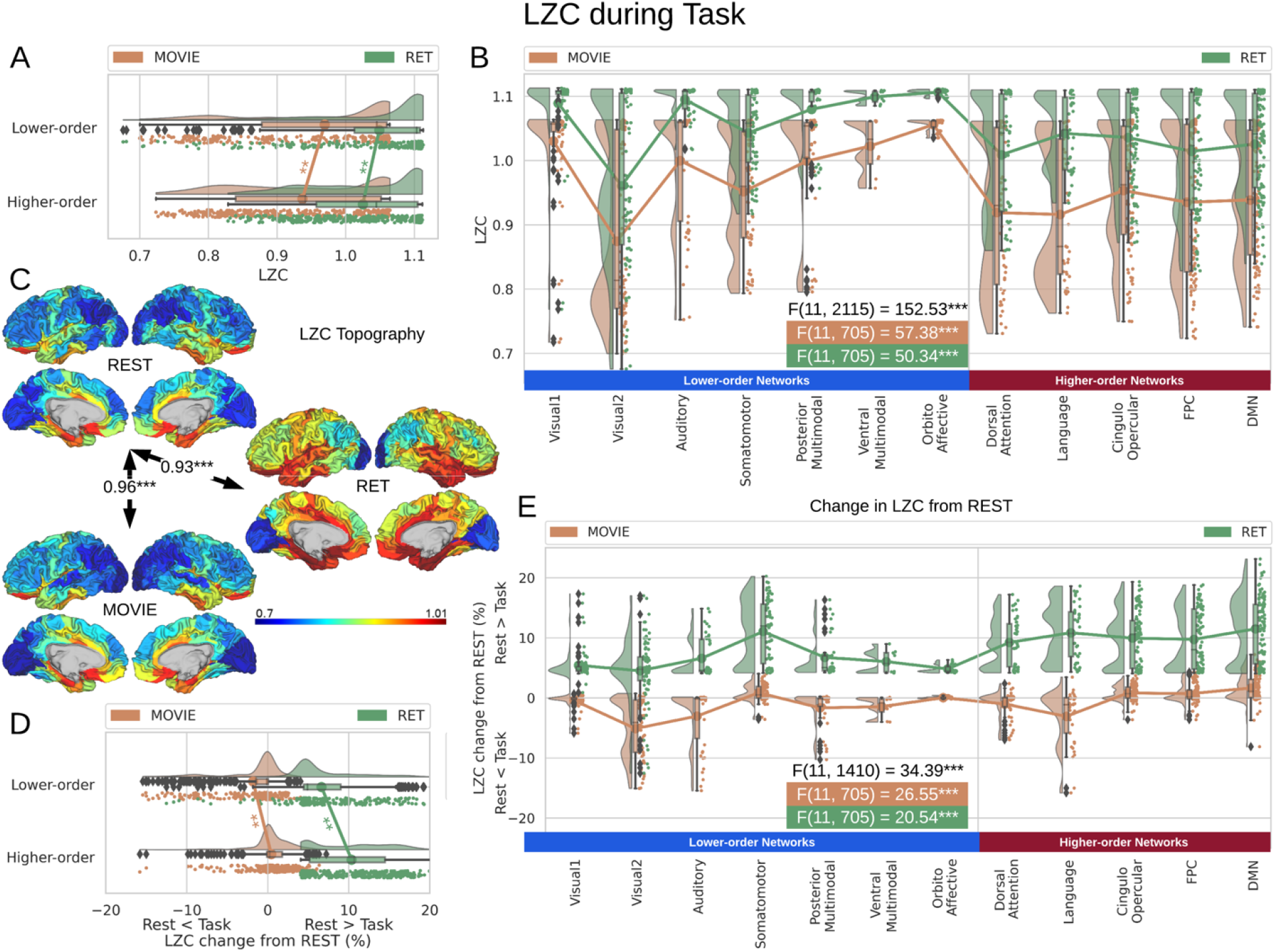
LZC during the task states. Points in the rain plots represent regions. **(A)** LZC during MOVIE and RET for lower- and higher-order networks. The paired student’s t-test between lower- and higher-order networks is significant with for MOVIE and for RET. **(B)** LZC during the two task conditions for the 12 networks. Three separate ANOVA models (One over the whole data, one for only the MOVIE, and one for only the RET data) showed significant differences among the networks. **(C)** Spatial distribution of LZC during REST, MOVIE and RET conditions alongside their corrected spatial correlation coefficients. The correlation is calculated over the regions to show the spatial similarity between resting and task states. **(D)** Change in LZC from REST to both task conditions for lower- and higher-order networks. Similar to task state results, the change was also significantly different between the two categories. **(E)** Change in LZC from REST to MOVIE and RET is also significantly different among the 12 networks. Stars represent the significance level ().

Moreover, dividing the LZC values into the 12 networks (Fig. 3B) and conducting similar twoway ANOVA (factors: networks with 12 levels and task with 2 levels) on them also showed significant (*p* < 0.001) effect of the network on LZC values (*F*(11, 2115) = 152.53, *η*^2^ = 0.32). Next, two separate one-way ANOVAs were designed to further test the effect of the network on LZC in each task. Both models showed significant *p* < 0.001 differences among the networks in LZC values (MOVIE: *F*(11, 705) = 57.38, *η*^2^ = 0.47; RET: *F*(11, 705) = 50.34, *η*^2^ = 0.43).

The differences in LZC between lower- and higher-order categories and also among the networks in the tasks were analogous to the previous resting state results, suggesting that the topography of LZC during task states was similar to the resting state. To test this hypothesis, we, in a second analysis, calculated the spatial correlation between the resting state (REST) and each task condition (MOVIE and RET, Fig. 3C). For each pair of conditions, LZC values were first averaged across subjects yielding a pair of LZC maps, then the two maps were correlated over regions, using Pearson correlation, yielding a single correlation value. The result showed significantly high correlation coefficients for both MOVIE (*α* = 0.96, *p* < 0.001) and RET (*α* = 0.93, *p* < 0.001). These data, further support the assumption that the resting state LZC topography is preserved during the two tasks.

Even though resting state LZC topography was similar in resting and task states, we, nevertheless, observed increases and decreases in LZC during the two task states. Therefore, in a third step, we calculated the percent of LZC change from the resting state for each region. For each task, the regional LZC values were subtracted from and then divided by their corresponding resting state values ((resting state − task state)/resting state); thus, normalizing the task state against the resting state, i.e., rest-task difference. We used the statistical models we applied during the task states, also to rest-task differences, i.e., to their percentage change values. Twoway ANOVA with network order (lower vs. higher) and task condition (MOVIE vs. RET) showed significant (*p*< 0.001) effect of network order on the percentage of change (*F*(1, 1430) = 173.06, *η*^2^ = 0.04), and further student’s t-test showed a higher percentage of change in higher-order networks compared to lower-order ones in both MOVIE (*t* = 7.28, *d* = 0.54, *p* < 0.01) and RET (t = 12.04, *d* = 0.90, *p* < 0.01). On the network level, similar to task analysis, significant (*F*(11, 1410) = 34.39, *η*^2^ = 0.07, *p* < 0.001) effect of the network in a twoway ANOVA model (network and condition) was followed by significant differences among the networks in two separate one-way ANOVAs for MOVIE (*F*(11, 705) = 26.55, *η*^2^ = 0.29, *p* < 0.001) and RET (*F*(11,705) = 20.54, *η*^2^ = 0.24, *p* < 001).

Although percentage of LZC change was statistically different between lower- and higher-order categories (Fig. 3D), and over the 12 networks (Fig. 3E), it presented distinctive results for the two tasks. In the MOVIE, positive values (decrease in LZC from REST to MOVIE) were observed in visual, auditory, and language networks. In contrast, LZC values were prominently increased from REST to our second task, i.e., RET (negative percentage of change) in all networks. Taken together, LZC topography during rest along the lines of lower- and higher-order networks is largely preserved during and thus carried over to the different task states. Additionally, one can nevertheless observe task-related changes in LZC once one subtracts task from rest; these concerned LZC differences between lower- and higher-order networks during the tasks. Hence, task-related LZC changes seem to loosely reflect the different input structures of our two tasks, which further supports the assumption that ISF do indeed carry information.

### Temporal dynamics - Median Frequency in rest and task states

To link the LZC in resting and task states to the ISF themselves, we characterized the distribution of their power spectral density (PSD) using the Median frequency (MF). MF is defined as the frequency which divides the area under the power spectral density into two halves. It was chosen to provide a summary measure of the distribution of low versus high frequencies in the PSD of ISF (Fig. 4A, see also Methods for the details).

**Fig. 4.**
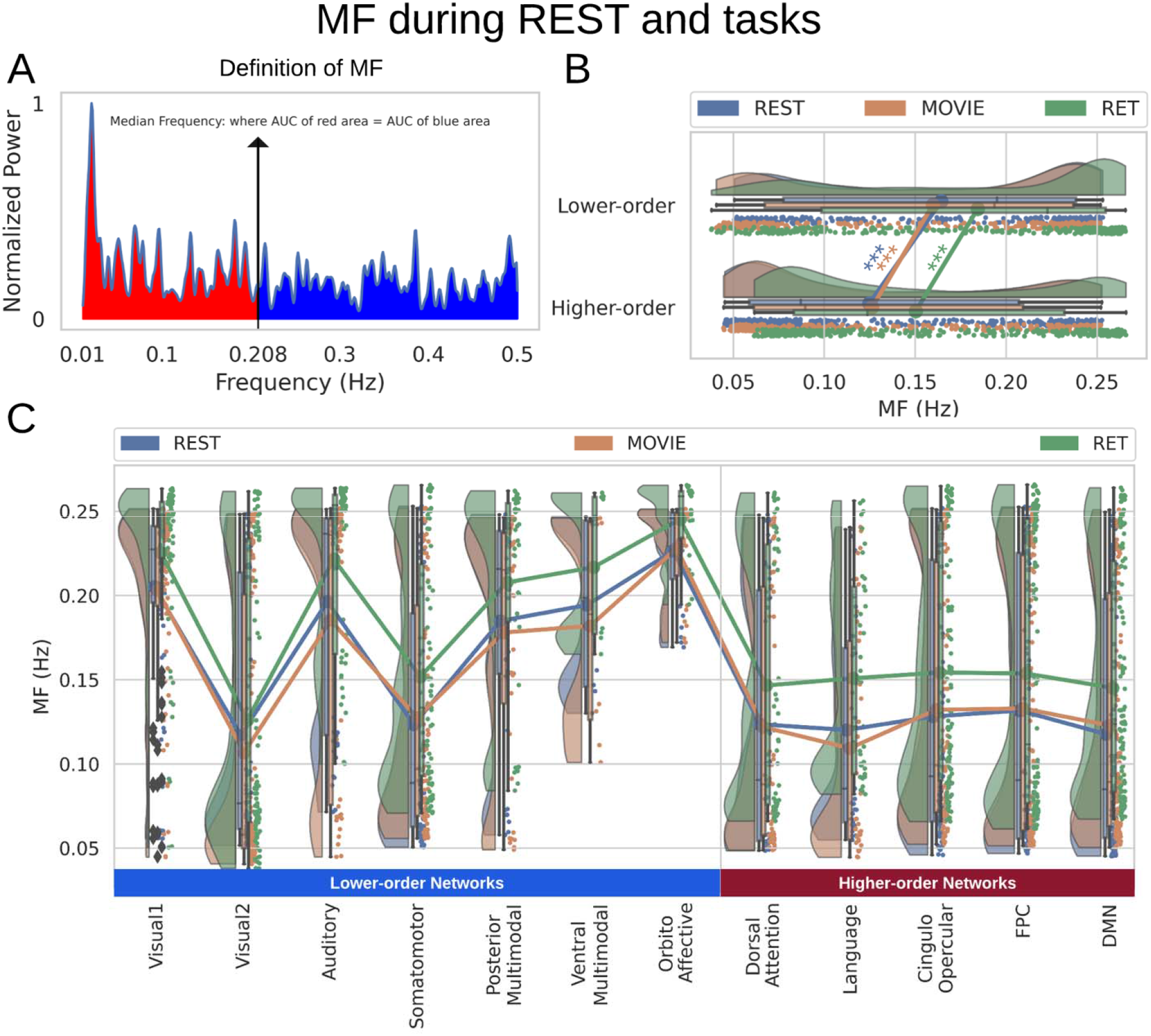
MF during both resting and task states. Points in the rain clouds represent regions. Stars represent the significance level (). **(A)** A power spectrum of a sample signal. Median frequency is calculated as the frequency at which the area under the curve (AUC) of power up to that frequency (red area) is equal to AUC beyond that point (blue area). **(B)** Similar to LZC, MF is significantly higher in lower-order networks compared to higher-order ones. **(C)** MF is also significantly different among the 12 networks during both resting and task states.

Calculating MF over different brain networks revealed a topographical distribution analogous to the one of LZC (Fig. 4B and Fig. 4C). Student’s t-test showed significantly () higher MF in lower-order networks compared to higher-order ones in both resting () and task states (MOVIE:; RET:). Furthermore, tested with one-way ANOVA, the topographical distribution was also significantly (different among the 12 networks (REST:; MOVIE:; RET:). These results are also replicated in the 3T dataset (Supp. Fig. 2). Together, these results show that the topographical differentiation of lower- and higher-order networks also holds on temporal grounds, namely based on the power spectral density of the ISF as measured by MF.

### The relationship between LZC and MF

The similarity in the topographical distribution of LZC and MF raises the question of their relationship. Specifically, it allows us to address our second key questions, namely “how” ISF process information. This was addressed in four steps. First, for each condition (REST, MOVIE, or RET) we calculated the regional correlation between LZC and MF. Regional correlation is calculated per region across subjects as the Pearson correlation between LZC and MF values (Fig. 5A). The p-values were corrected for multiple comparisons using the False Discovery Rate (FDR) method at *α* = 0.05 while non-significant coefficients were ignored. This revealed a variety of correlation values for the LZC-MF relationship across the wide range of 0.43 to 0.88 across the different regions. Given such a wide range of correlation values, we assumed that topographical differences strongly shape the LZC-MF relationship; this was addressed in subsequent analyses.

**Fig. 5.**
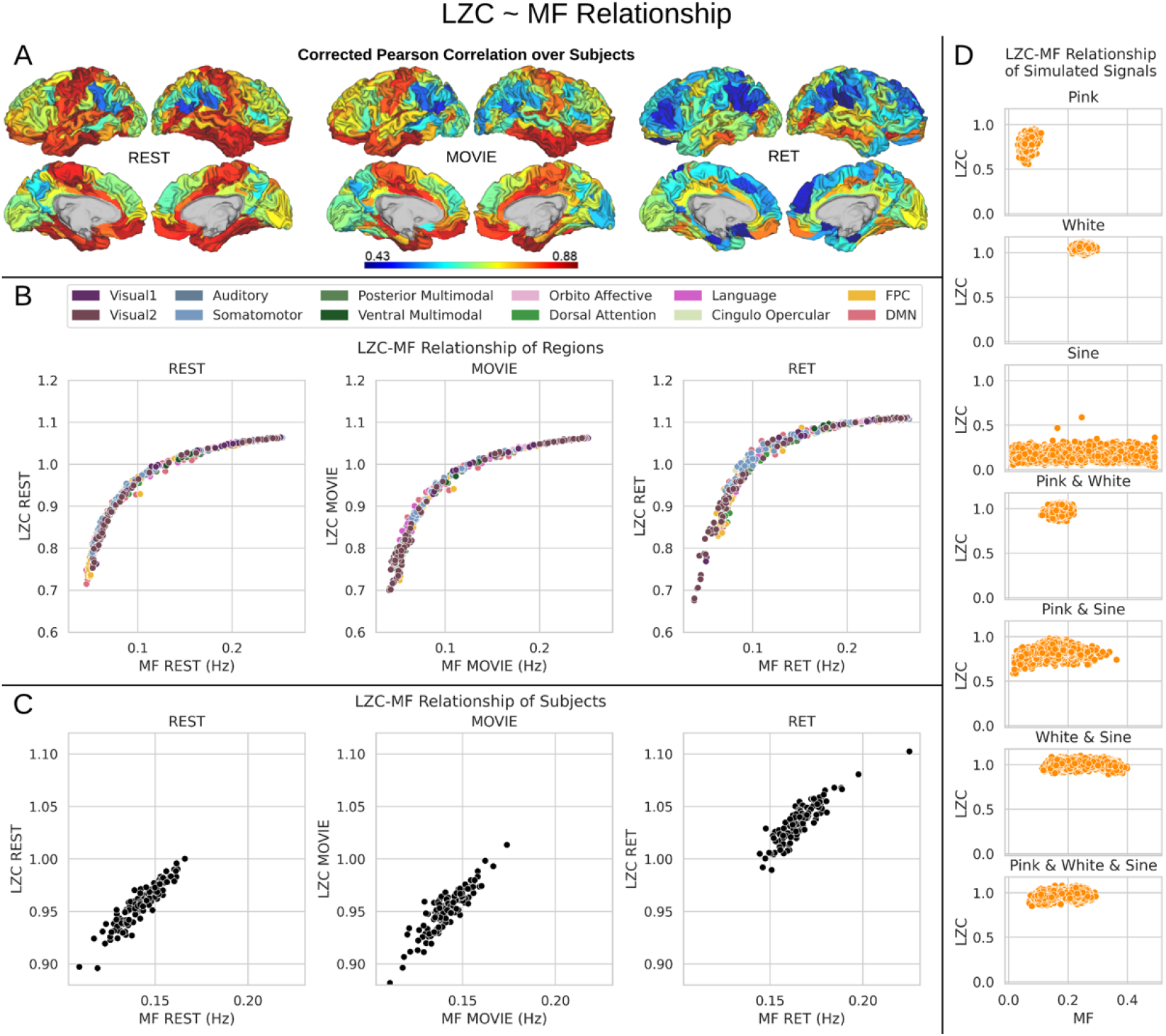
The relationship between LZC and MF. **(A)** Regional Pearson correlation between LZC and MF for the three conditions of REST, MOVIE and RET. The p-values are corrected using the FDR method at. The wide range of correlation values (0.43 - 0.88) suggests topographical differences affect the LZC-MF relationship. **(B)** Regional scatter plots of LZC-MF relationship for the three conditions. Each point is a region averaged over subjects, and the colors show the regions belonging to specific networks. All plots show LZC as a nonlinear function of MF. **(C)** Scatter plots of LZC-MF relationship for different subjects in the three conditions. Each point is a subject averaged over regions. The nonlinear relationship can no longer be observed. **(D)** Simulation of the LZC-MF relationship with seven different types of signal. Each plot shows the distribution of LZC as a function of MF for 5000 simulated signals. The nonlinear relationship cannot be observed in the simulated signals suggesting that it is unique to the brain signal.

In the second step, the LZC-MF relationship was explored in more detail by looking at their values in different brain regions. First, both LZC and MF values were averaged over subjects and then each region’s LZC value was plotted against its corresponding MF value (Fig. 5B). This suggested a non-linear regime between LZC and MF over brain regions across all three conditions. Specifically, we observed that regions with lower MF in rest exhibit largely different LZC values in both rest and task whereas regions with higher MF in rest no longer showed marked LZC differences in both rest and task. Together, this amounts to a non-linear relationship in the topography of LZC and MF with high and low MF exerting different impacts on LZC (also replicated in 3T dataset in Supp. Fig. 3).

We tested whether this non-linear relationship is specifically related to the topographical distribution of LZC and MF (topographical relationship) rather than inter-individual differences between subjects (Step 3). To do so, LZC and MF values were averaged over all regions leaving a pair of MF and LZC values per subject (subject-based relationship). This (Fig. 5C) revealed a relationship different from the topographical one, suggesting a linear trend for the subject-based relationship of LZC and MF: the higher the MF in a specific individual (across all its regions), the higher its LZC (across all its regions, Fig. 5C). One caveat is that the range of value for the calculation of inter-individual LZC-MF relationship is lower than the one in our regional topographic LZC-MF; for that reason, we cannot fully exclude potential non-linear regime in inter-individual LZC-MF relationship.

As the fourth step, the LZC-MF relationship was further explored by using the brain’s *T*_1_*w*/*T*_2_*w* values (see Methods) to test whether it is structurally based. *T*_1_*w*/*T*_2_*w* is suggested to provide a non-invasive proxy of anatomical hierarchy in primate cortex ^66^. For both LZC and MF, each region’s values were averaged over subjects and then the regional distribution was plotted as functions of values (Supp. Fig. 4A for LZC and B for MF). This also failed to show the specific topographical relationship between LZC and MF observed previously. That, suggests the non-linear relationship of LZC and MF to hold independent of their underlying anatomical relationship; we thus assume that the non-linear topographical LZC-MF relationship is primarily driven by the dynamics of ISF, rather than being based on structural anatomical grounds (also replicated in 3T dataset in Supp. Fig. 3).

### Simulation of the relationship between LZC and MF

As a final confirmatory analysis to show the truly topographical nature of the non-linear LZC-MF relationship, we decided to analyze simulated data. Seven different categories of pseudo-aleatory signals (without any topographical distribution) were simulated in the same frequency range (0.01-0.5) as our data with the same sampling rate. The categories were pink noise, white noise, sine wave, and their linear combinations (e.g. white and pink noises or white noise and sine wave). Each category contained 5000 signals (total of 35000, see Supp. Fig. 5 for a sample signal in each category) with different combinations of power set randomly in the same ranges as the real signals.

Pink noise was utilized to simulate scale-freeness ^16^, sine wave for oscillation and white noise for pure randomness. The parameters for each signal and the weights for their linear combinations used random values chosen from uniform distributions to bound the signals in the predefined frequency range of our real data. Calculating LZC and MF on these signals and plotting them against each other (Fig. 5D) showed no specific relationship between the two measurements. These results, thus, suggest that the non-linear relationship between MF and LZC is related to the topographical distribution of low and high MF values in different brain regions rather than remaining independent of such topographical distribution. These results also provide evidence for the dissociation between LZC and MF, as the correlation between them is different depending on the signals assessed. Moreover, our simulation shows that LZC is not measuring noise in the BOLD signal of fMRI.

### The relationship between LZC change from resting state and MF during resting state: the dynamic range of LZC

Is the change of LZC from rest to task in specific regions dependent upon the power spectral density, i.e., MF, of that particular region? Answering this question will allow us to address our second key question, namely “how” ISF process information. If rest MF mediates LZC rest-task differences, one would strongly assume that the power spectral density (MF) of ISF processes changes in information load during the transition from resting to task state as measured by LZC. We, therefore, investigated how the degree of change in LZC from resting to task state is related to the power spectrum during rest (MF-REST). The “change” values of all regions were plotted against their corresponding MF values during rest (MF-REST) for both task conditions (Fig. 6A). Careful consideration of these plots alongside a mediation analysis of LZC with MF as a mediator (see supplementary results for details) suggested that there is a relationship between the range of change in LZC values and MF-REST.

**Fig. 6.**
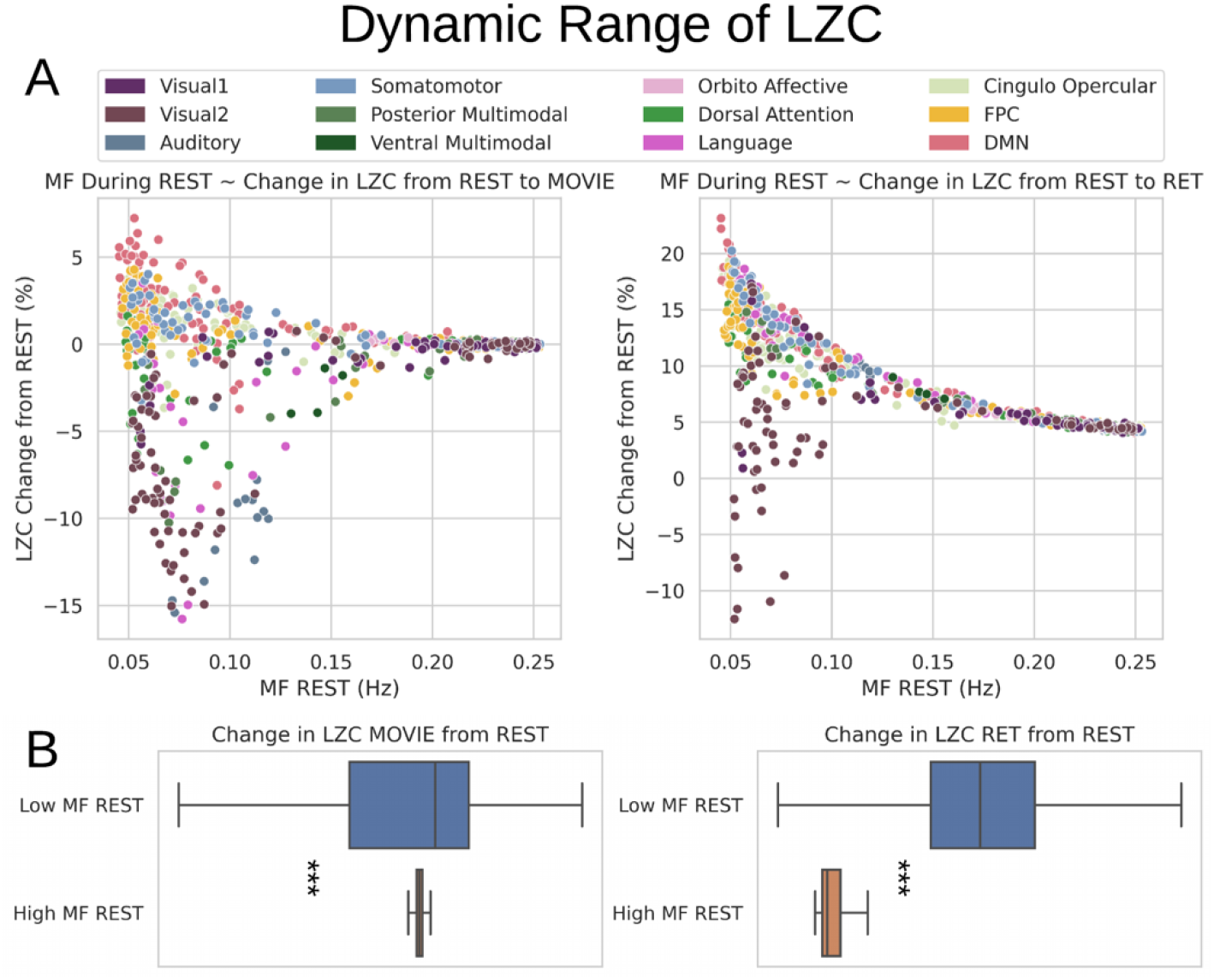
The relationship between the change in LZC from REST and MF during REST. **(A)** The scatter plot of MF-REST (x-axis) vs. the change in LZC (y-axis) from REST to MOVIE (left) and RET (right). The plots indicate a decrease in the range of the “change” as a function of MF-REST. **(B)** Box plots showing the range of the “change ” values for low-MF and high-MF during REST. MF-REST was median-split into low and high categories. Statistics were conducted using student’s t-test over subjects. For each subject, a range of LZC change was calculated in low and high MF categories. For both conditions (MOVIE and RET) the range of LZC change in low-MF is significantly wider than high-MF. Stars represent the significance level ().

Following that, we decided to conduct a median-split of the MF-REST values across brain regions. That served the purpose of calculating the “change” values in dependence on low MF-REST and high MF-REST. Plotting both categories with box plots (Fig. 6B), revealed a large difference in the range of change in LZC values (i.e. the range of change in LZC from resting to task states) between low and high MF-REST categories. To test whether this difference is significant or not, the “change” values of each subject were divided into two categories of low and high MF-REST. Then, the significant () difference between the two was confirmed by Student’s t-test over subjects (MOVIE:; RET:).

These data suggest, that the range of LZC change values in low MF-REST regions **is** significantly higher than in high MF-REST regions. Together, these results suggest that the range in which the region’s LZC can change is associated with the level of MF-REST (low or high). Therefore, we speak of a dynamic range of LZC which we hypothesize to be related to PSD of the region as indexed by MF: low rest MF yields a large dynamic range of LZC, i.e., a wide range of rest-task differences among its regions, while high rest MF is related to small a dynamic range of LZC, i.e., narrow range of rest-task differences among its regions.

To further test the dynamic range of LZC, we incorporated a moderation model to see if MF-REST being low or high can moderate the relation between LZC during rest and task states. A binary moderator variable (Z) for MF-REST being low (Z = 0) or high (Z = 1) was defined. Then, the effect of LZC-REST and Z was explored on LZC during the two task conditions using linear regression (see Methods for details). The model showed significant moderation of the LZC rest-task relationship with MF-REST being low or high. For MOVIE the effect of the moderator was 0.08 (*t* = 9.63, *p* < 0.001). Likewise, for RET the effect of the moderator was 0.42 (*t* = 44.52, *p* < 0.001). In other words, our moderation analysis suggests that the effect of LZC during REST on LZC during MOVIE (or RET) is moderated by MF during REST.

Taken together, these results show strong evidence for the relationship of MF and LZC in a non-linear topographical way. The level of MF in a region’s resting state drives its propensity for change in its information processing during the transition from resting to task state as measured by LZC. Regions with low rest MF indicating strong power in the very slow ranges of ISF can yield a larger spectrum of LZC changes during the task (relative to rest) than regions with high MF. Hence, the very slow frequency ranges of resting state MF seem to foster larger dynamic ranges in the regions’ information processing as manifest in rest-task differences. This addresses our second key question, namely “how” ISF process information: they process information through their power spectral density as indexing their “velocity” or “speed”.

### Replication with 3T data

#### Topographical resting state LZC/MF of 3T data

3T resting state data was analyzed using the same procedure as our 7T data (Supp. Fig. 1 for LZC and Supp. Fig. 2 for MF). The difference between lower- and higher-order networks was statistically tested with student’s t-test between the two across regions which showed that both LZC and MF is significantly (*p* < 0.001) higher (LZC: *t* = 3.60, *d* = 0.27; MF: t = 5.30, *d* = 0.40) in lower-order networks compared to higher-order ones. Furthermore, LZC and MF were also significantly (*p* < 0.001) different among the 12 networks tested with one-way ANOVA (LZC: F(11,705) = 44.15, *η*^2^ = 0.40; MF: F(11, 705) = 52.22, *η*^2^ = 0.44). These results validate our original topographical findings.

#### The relation between LZC and MF in 3T data

As another confirmatory analysis, the relationship between LZC and MF was investigated in resting state 3T data. Like before, first the two measurements were correlated with each other using Pearson methods (Supp. Fig. 3A). Moreover, LZC was plotted as a function of MF both for each region (Supp. Fig. 3B left) and for each subject (Supp. Fig. 3B right). The non-linear relationship over the brain regions was again observed in the 3T data. Both regional (averaged over subjects) LZC and MF were separately plotted as functions of *T*_1_*w*/*T*_2_*w* values which didn’t show any specific non-linear relationship (Supp. Fig. 3C). Taken together, these results replicate our findings on LZC and MF relationship.

## Discussion

We here investigate the role of ISF in information processing by addressing “whether” and “how” ISF carry and process information in resting and task state fMRI. Our main findings in both main (7T HCP) and replication (3T HCP) data sets are: (I) significantly different resting state LZC and MF in lower- and higher-order networks with the latter showing lower LZC and MF values; (II) task-related LZC changes during different tasks in lower- and higher-order networks; (III) non-linear topographical relationship of LZC and MF with regions showing lower rest-MF values being related to lower resting state LZC and larger capacity for task-rest changes in LZC. Together, these findings provide evidence that ISF carry and process information (LZC) through their power spectral density, i.e., ‘velocity’ or ‘speed’ (MF).

### Do ISF carry information? Topographical differences in complexity and dynamics of lower- and higher-order networks

The first key question is “whether” ISF carry information. Our findings strongly suggest that this is indeed the case. We show different degrees of LZC in different networks during the resting state. Higher-order networks like the DMN, FPN, language, and dorsal attention exhibit lower values of LZC in the resting state. In contrast, lower-order networks like sensory networks, orbito affective, and multimodal show high LZC values. Hence, taken together, the ISF signal is more regular and more compressible, i.e., lower LZC, in higher-order networks than in lower-order networks. More generally, this means that lower-order networks carry more information in their signal, i.e., it cannot be as much compressed as measured by LZC, than higher-order networks with their lower LZC.

Our findings complement previous results that show analogous differences between lower- and higher-order networks. Spatially, lower-order networks are more locally connected while higher-order networks are more globally connected throughout the whole brain. This led, recently, to the so-called core-periphery organization ^26,28^: Lower-order unimodal networks like sensory networks are at the periphery while higher-order transmodal networks like DMN and FPN constitute the core as they are connected throughout the whole brain. Recent data suggest such core-periphery organization to be mirrored also in the brain’s temporal organization as its intrinsic neural timescales are shorter in the lower-order networks, i.e., the periphery, and longer in the higher-order networks, i.e., the core ^58–60,67–69^. Our data extend these findings by showing that lower- and higher-order networks carry different degrees of information (LZC) which converges with corresponding differences in their temporal dynamics (MF). This, suggests that intrinsic information processing (LZC) and brain dynamics (MF) converge in the topographical differentiation of lower- and higher-order networks.

The differential role of lower- and higher-order regions in information processing is further supported by our results in task states. We observed network- and task-specific changes during the two tasks relative to the resting state. Together, extending previous findings ^42^, the observed task-rest differences in LZC further support the assumption of task- and network-specific changes in LZC and, more generally, the involvement of ISF in carrying information in both resting and task states.

### How do ISF process information? ISF process information (LZC) through their temporal dynamics (MF)

So far, we have addressed the question of “whether” ISF carry information. That was addressed by demonstrating that the ISF show topographical differences in LZC during rest and task which follows the well-established information hierarchy of lower- and higher-order networks. This leaves open “how” the ISF process information, that is, through what underlying mechanism? To address this question, we included median frequency (MF) as a measure of brain dynamics operating by the characterization of the PSD distribution. We saw a similar topographical pattern in MF as in LZC. This suggests a close relationship of LZC and MF meaning that the power spectral density of ISF may mediate and thus process their information. Probing this assumption, we tested various correlation and simulation models.

Spatial correlation confirmed such a close relationship; however, regional MF-LZC correlation revealed quite a variety of different degrees in their correlation within the different regions. Such a wide variety of LZC-MF correlations across different regions suggests a differential relationship of regions with low and high MF to LZC. This was confirmed by subsequent analyses. We obtained a non-linear topographical pattern in the LZC-MF relationship in both rest and task states. Regions with low MF at rest showed lower rest LZC and more differentiation in their respective LZC rest-task changes than those regions exhibiting high rest MF. We could further demonstrate that such non-linear relationship is related specifically to the topographical distribution of low and high MF values rather than being associated with inter-individual differences, anatomical differences in *T*_1_*w*/*T*_2_*w* of the different regions, or some other kinds of LZC-MF relationship independent of their topographical distribution (as tested for in simulation).

Further, we observe that the MF is related to the range of possible LZC rest-task changes. Regions with low MF values during rest show much larger changes in their rest-task LZC values than those with high MF (see supplementary results). Hence, MF in rest seems to modulate the capacity for change in LZC during the transition from rest to task: regions with low MF and thus more power in the slower ISF ranges have a higher likelihood of exhibiting larger changes in their information processing, i.e., increases or decreases, during the transition from rest to task states. Hence, it is the brain dynamics of the different regions’ resting state rather than the regions themselves (independent of their dynamics) that is related to the topographical distribution of LZC during rest and task.

How can brain dynamics (MF) during rest modulate and shape information processing (LZC) in ISF? Low values in MF reflect stronger power in the very infraslow frequency fluctuations which can be characterized by extremely long cycle durations. These long cycle durations are assumed to be ideal for summing and pooling different stimuli occurring at different points in time such that their respective information is processed and thus lumped together ^1,4,6,70,71^. Such summing and pooling may thus enable higher degrees of integration of temporally distinct stimuli. That, in turn, leads to less complex signal dynamics with lower LZC values. One would consequently expect that regions showing lower values in MF should also exhibit lower LZC values which is exactly what we observed in our data.

This supposed modulation of the LZC-MF relationship by the pooling and summing of stimuli through cycle duration may also account for the observed differences in the rest-task changes of the LZC in regions with low and high rest MF. Given the inverse relationship between power and frequency, i.e. the scale-freeness of the brain oscillations ^16^, if a region shows low MF, the power of its long cycle duration is stronger than the one of a region with higher MF. That may allow the low rest MF region to pool, sum, and ultimately integrate more external stimuli during task states ^70^ than a region with high rest MF. This, in turn, changes the degree of that region’s complexity and thus its LZC to a large degree. This stands in contrast to a high rest MF region that, due to its lower power in the long cycle duration, cannot pool and sum as many external stimuli and consequently cannot as much change its LZC from rest to task. That remains to be tested in future modelling studies though.

## Limitations

There are a few considerations that should be considered in this study. First, task-unspecificity should also be replicated using different tasks in different modalities and domains; however, concerning our metrics, we included two tasks with completely different complexity and temporal structure. The movie presents a continuous task while the retinopathy is an event-related trial-based discontinuous task. The inclusion of tasks with two different structures accounts for the recent suggestion ^72^ of considering and including tasks with different structures, i.e., continuous vs. trial-based. Second, this study contains no behavioral measurement, but on the other hand, the tasks measured pure perception and stimulus processing with no interference of cognitive demands, thus no-report paradigms as distinguished from report paradigms ^73^. The reliance on no-report paradigms allowed to isolate stimulus-related effects, i.e., movie and visual stimuli during retinopathy, as they are supposed to be related to, specifically, primary sensory networks like visual and auditory networks. Hence, the no-report paradigms are ideal to test the response of primary networks independent of any task-related confounds as in report paradigms.

## Conclusion

The infraslow frequency fluctuations (ISF) are more powerful than faster frequencies but less well understood. Here we investigate “whether” they carry information (LZC) and, if so, “how” they mediate information processing, namely through which of their features. In addressing the first question, our findings strongly support that ISF carry information as the LZC shows differences in lower- and higher-order networks in rest and task states as well as task-specific and -unspecific changes. Moreover, responding to the second question, we demonstrate that the degree of LZC rest-task changes is driven by rest MF levels in a non-linear topographical way. This suggests that ISF carry and process information (LZC) through their temporal dynamics, i.e., power spectral density as an index of their ‘velocity’ or ‘speed’. In conclusion, our findings demonstrate that ISF carry and process information through their temporal dynamics supporting 74,75 the recently introduced concept of “Spatiotemporal Neuroscience”^74,75^.

## Acknowledgments

This research has received funding from the European Union’s Horizon 2020 Framework Program for Research and Innovation under the Specific Grant Agreement No. 785907 (Human Brain Project SGA2). GN is grateful for funding provided by UMRF, uOBMRI, CIHR and PSI. We are also grateful to Chris J. Honey for giving useful suggestions. We are also grateful to CIHR, NSERC, and SHERRC for supporting our tri-council grant from the Canada-UK Artificial Intelligence (AI) Initiative ‘The self as agent-environment nexus: crossing disciplinary boundaries to help human selves and anticipate artificial selves’ (ES/T01279X/1) (together with Karl J. Friston from the UK)

## Materials and Methods

### Experimental Design

Data were selected from Human Connectome Project’s 7T dataset (rather than HCP 3T) to achieve a high signal-to-noise ratio (SNR) (and to have more fine-grained resolution) which is especially relevant for properly capturing temporospatial dynamics. Other reasons for choosing the 7T data were the longer and more continuous scanning during task states. We used HCP 3T resting-state data for validation purposes; in contrast, we did not use the task data of the 3T dataset because of their short block design which makes proper measurement of the dynamics with our measures impossible.

From the data of 1200 subjects released in HCP-1200 ^76^, 146 of them had completed the full 7T imaging protocol of the human connectome project (HCP) without any imaging quality issues. A total of 14 fMRI runs (TR=1s) including four resting-state (REST), four movie-watching (MOVIE), and six retinotopy (RET) of these participants were used in this study. Full details on data acquisition and preprocessing are provided in separate articles ^77,78^. In each MOVIE run, subjects had to watch a movie of approximately 15 minutes consisting of several clips separated by 20-seconds rest periods. Different clips were used in different runs (details are available at HCP S1200 Release Reference). For RET, stimuli were constructed by creating slowly moving apertures and placing a dynamic colorful texture within the apertures [Check ref. for details].

Preprocessed data was downloaded from the HCP database at https://db.humanconnectome.org. The preprocessing pipeline includes registration to MNI space, alignment for motion, fieldmap correction, FIX denoising, and MSMAll group registration. Full details of all the steps are available in ref. ^77,80^.

### Data Processing & Statistical Analysis

All steps of the data processing were performed with in-house scripts written in Python programming language using numpy, scipy, cifti, joblib, matplotlib, and seaborn libraries. The source code is freely available at www.georgnorthoff.com/code/. For brain map visualization purposes, wb_view (part of Connectome Workbench software) was used.

All statistical analyses were performed in the statsmodel library ^81^ in Python and R v.3.6 and all p-values were corrected for multiple comparisons using the False Discovery Rate (FDR) method. The student’s t-test was used to measure the statistical differences between lower- and higher-order networks. Moreover, Analysis of Variance (ANOVA) ANOVA was used to investigate the difference between the networks during each condition.

#### Template and network definition

The preprocessed data were high-pass filtered at 0.01 to both maintain the full frequency spectrum of the data ^82^ and remove noise. After that, signals were averaged over brain voxels defined in the template provided by ref. ^62^ to get one fMRI signal per region per subject per run. The template contains 717 regions categorized into 12 networks of visual1, visual2, auditory, somatomotor, posterior multimodal, ventral multimodal, orbito affective, dorsal attention, language, cingulo opercular, frontoparietal, and default-mode. To investigate our lower-order vs. higher-order hypothesis, we also divided the regions into lower-order and higher-order (HON) networks. Lower-order networks included the regions in the visual1, visual2, auditory, somatomotor, posterior multimodal, ventral multimodal and orbito affective networks. Regions inside dorsal attention, language, cingulo opercular, frontoparietal, and default-mode networks were put under the higher-order networks category.

#### Calculation of Lempel-Ziv complexity

To calculate the Lempel-Ziv complexity (LZC, Fig. 1 white box), each signal was converted into a binary sequence. Previous studies ^39,64^ suggest that the median of the signal’s amplitude is a good candidate to use as a binarization threshold. After binarizing each region’s signal and converting it to a string sequence, the Lempel-Ziv algorithm ^38^ was used to compute LZC. As earlier studies have pointed out, LZC is dependent on the length of the sequence, thus a normalization factor (more info in ^39^) was used to remove that effect. LZC processing is illustrated in Fig. 1A.

#### Calculation of Median Frequency

Median frequency (MF) was measured by first calculating the power spectral density (PSD) of each region’s signal. PSD was calculated using the Welch algorithm ^83^ with a Hanning window implemented in the Scipy package of Python programming language. The median frequency (MF) was calculated from the PSD as the frequency which divides the area under the curve of OSD into two halves (Fig. 4A).

#### Resting and task state topographical calculations

The values of both measurements (i.e. LZC and MF) were first calculated for each of the 14 fMRI runs and then averaged over the runs in all three conditions (REST, MOVIE and RET). After that, based on the requirements of each analysis the values were either averaged over the lower- and higher-order division or the 12 aforementioned networks. The lower-order category was compared to the higher-order using student’s t-test over regions.

#### Rest-task similarity

The similarity between resting and task states was addressed using spatial and regional correlations. The spatial correlation was calculated as a single Pearson correlation coefficient between resting and a task condition (e.g. REST vs. MOVIE) over brain regions after averaging over subjects (thus creating a single brain per condition). The regional correlation was used to measure the regional s9imilarity between resting and task states. This correlation was calculated for each region across subjects between a pair of conditions (e.g. LZC during REST and MF during REST).

#### Percentage of change from resting state

The difference between resting and task states was calculated as a percentage of change per region per subject. Each region’s REST and task (e.g. MOVIE) values were put in the (REST − TASK) x 100/REST. formula to both measure the change from resting to task state and normalize against rest at the same time. Then, for each subject, the percentage of change values were either averaged over the 12 networks or along the lower- and higher order division. The difference between lower- and higher-order networks and the difference among the networks were statistically tested using the student’s t-test and ANOVA methods.

#### *T*_1_*w*/*T*_2_*w* map

The ratio of T1- to T2-weighted images is suggested to provide a non-invasive neuroimaging proxy of anatomical hierarchy in the primate cortex ^66^. So, we used it to investigate whether the relationship between LZC and MF is anatomically based. A cortical map containing the *T*_1_*w*/*T*_2_*w* was provided with the HCP dataset. The map was MSMAll registered and bias field-corrected. We parcellated the map into the regions/networks of our template by averaging the *T*_1_*w*/*T*_2_*w* values of voxels into different regions.

#### LZC-MF simulation

The relationship between LZC and MF was explored using simulated signals. 35000 pseudo-aleatory signals within seven different categories (5000 each, see Supp. Fig. 5 for a sample signal in each category) were simulated to investigate whether the LZC-MF relationship is non-linear or not and whether the relationship is specific to the brain. The seven categories were (I) pink noise, (II) white noise, (III) sine wave, and linear combinations of (IV) pink and white noises (V) pink noise and sine wave (VI) white noise and sine wave and (VII) pink noise, white noise and sine wave.

The weights of the signals linear combinations were chosen randomly. All random values were chosen from a uniform distribution and were controlled to produce signals in the same frequency range of our original data (0.01-0.5 Hz). Pink noise was chosen to model the scale-free behavior ^16^, white noise for pure randomness and sine wave for oscillation. The MF and LZC were calculated for each signal and used to further investigate the relationship between the two measurements, and validate LZC calculations in fMRI data.

#### Change in LZC from REST and MF-REST mediation model

To investigate the relationship between MF during resting state (MF-REST) and the change in LZC task from resting state, a mediation analysis was performed using the mediation library in R. Two separate models were created for the two task conditions. Each model consisted of MF-REST as the mediator, LZC during REST as the independent variable (IV) and LZC during a task condition as the dependent variable (DV). After running the model and observing the indirect effects of IV on DV, the significance of that effect was tested using bootstrapping procedures ^84,85^. Unstandardized indirect effects were computed for each of 1000 bootstrapped samples, and the 95% confidence interval was computed by determining the indirect effects at the 2.5th and 97.5th percentiles.

#### Change in LZC from REST and MF-REST moderation model

To further investigate the relationship between the change in LZC from REST and MF-REST, regions were divided into two categories based on their MF-REST values. The median of MF-REST was used as the threshold and the regions with lower (higher) MF-REST than the median were put into the low (high) MF-REST category. The median was used to balance the two categories. A new binary variable (Z) was created for each region’s MF-REST category (0 = low MF-REST and 1 = high MF-REST). Z was injected in a linear regression model as a moderator: *Y* = *β*_1_X + *β*_1_Z + *β*_1_XZ + *β*_4_. In the linear regression equation, Y is LZC during a task condition, X is LZC during resting state, and Z is the binary value of MF during resting state.

