## Supplementary Material for "The interplay between information flux and temporal dynamics in infraslow frequencies"

### Supplementary Results

#### Rest to task mediation analysis

To explore whether MF during rest has any effect on the change in LZC from rest to task state, we performed a mediation analysis on the LZC and MF-REST values (Supp. Fig. 6). In the model, the effect of LZC-REST on LZC-MOVIE (LZC-RET) was investigated considering MF-REST as the mediator. This allowed us to explore whether the relationship between LZC during REST and MOVIE (RET) is mediated by MF during REST. The indirect effect was 0.84 * 0.23 = 0.17 (-1.01 * 0.34 = -0.34). We tested the significance of this indirect effect using bootstrapping procedures ^1^ in the mediation library ^2^ of R. Unstandardized indirect effects were computed for each of 1000 bootstrapped samples, and the 95% confidence interval was computed by determining the indirect effects at the 2.5th and 97.5th percentiles.

The bootstrapped unstandardized indirect effect was 0.16 (-0.24), and the 95% confidence interval ranged from 0.10 to 0.22 (-0.29 to -0.19). Thus, the indirect effect was statistically significant (p < 0.001) showing that MF-REST can indeed be considered as a partial mediator for the change in LZC from rest to task states. It is worth to mention that the mediation analysis was performed with (the above results) and without averaging over subjects to control for the inter-individual effect in bootstrapping. In the with-averaging analysis, first, the data was averaged over subjects then fed into the model, but in the latter one, the subject-level data was used.

In the without-averaging model, the subject-level data was fed into our mediation model. Like before, LZC-REST was used as the independent variable, LZC-MOVIE (LZC-RET) as the dependent one and MF-REST as the mediator. The indirect effect was 0.64 * 0.33 = 0.21 (-0.64 * 0.14 = -0.08). The bootstrapped unstandardized indirect effect was 0.21 (-0.09), and the 95% confidence interval ranged from 0.21 to 0.22 (-0.099 to -0.09). Thus, the indirect effect was statistically significant (p<0.001), confirming our original with-averaging results.

### Supplementary Figures:

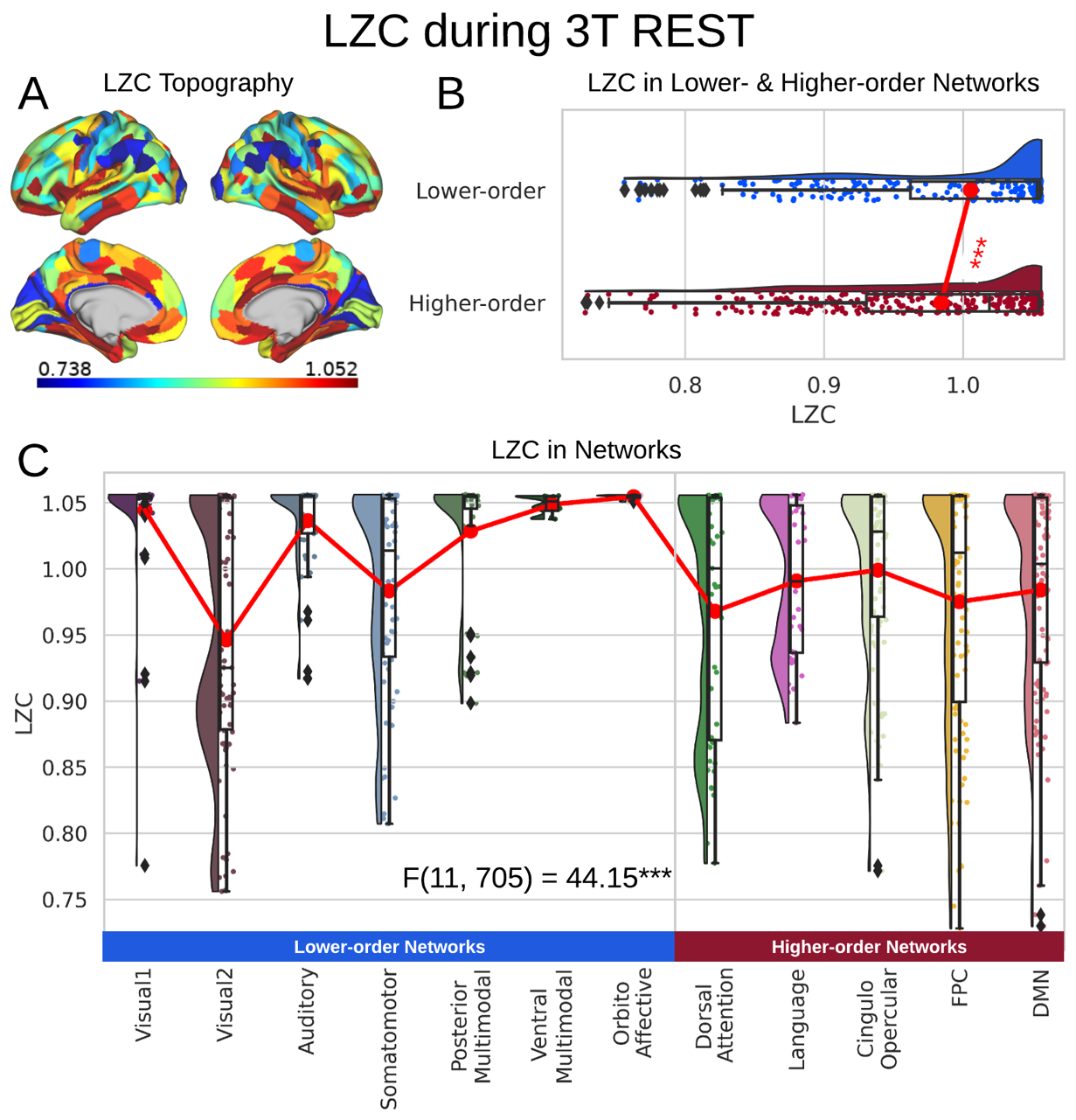

**Supp. Fig. 1. LZC during the 3T resting state.** Rainclouds represent regions. **(A)** Spatial distribution of LZC during 3T REST condition. **(B)** LZC during rest for lower- and higher-order networks. Student’s t-test shows significant ($p<0.001$) differences between lower- and higher-order networks ($t=3.60, d=0.27$). **(C)** LZC during REST for the 12 networks. One-way ANOVA showed a significant ($p<0.001$) difference among the networks ($F\left( 11, 705 \right)=44.15, \eta^{2}=0.40$). Stars represent the significance level ($*** \equiv\alpha=0.001$)

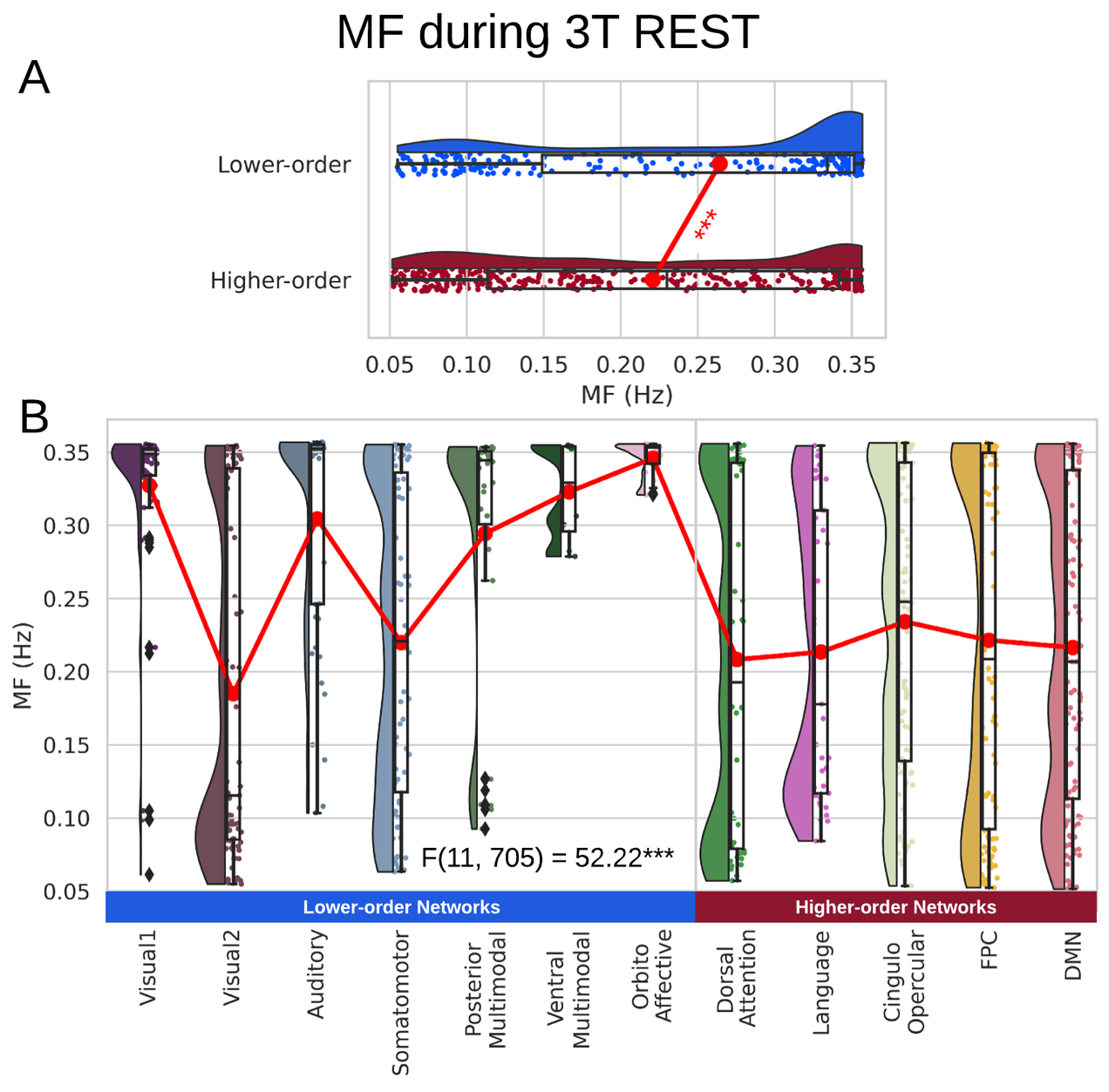

**Supp. Fig. 2. MF during the 3T resting state.** **(A)** MF during rest for lower- and higher-order networks showing higher MF in lower-order networks. **(B)** LZC during rest for the 12 networks. One-way ANOVA showed a significant difference among the networks.

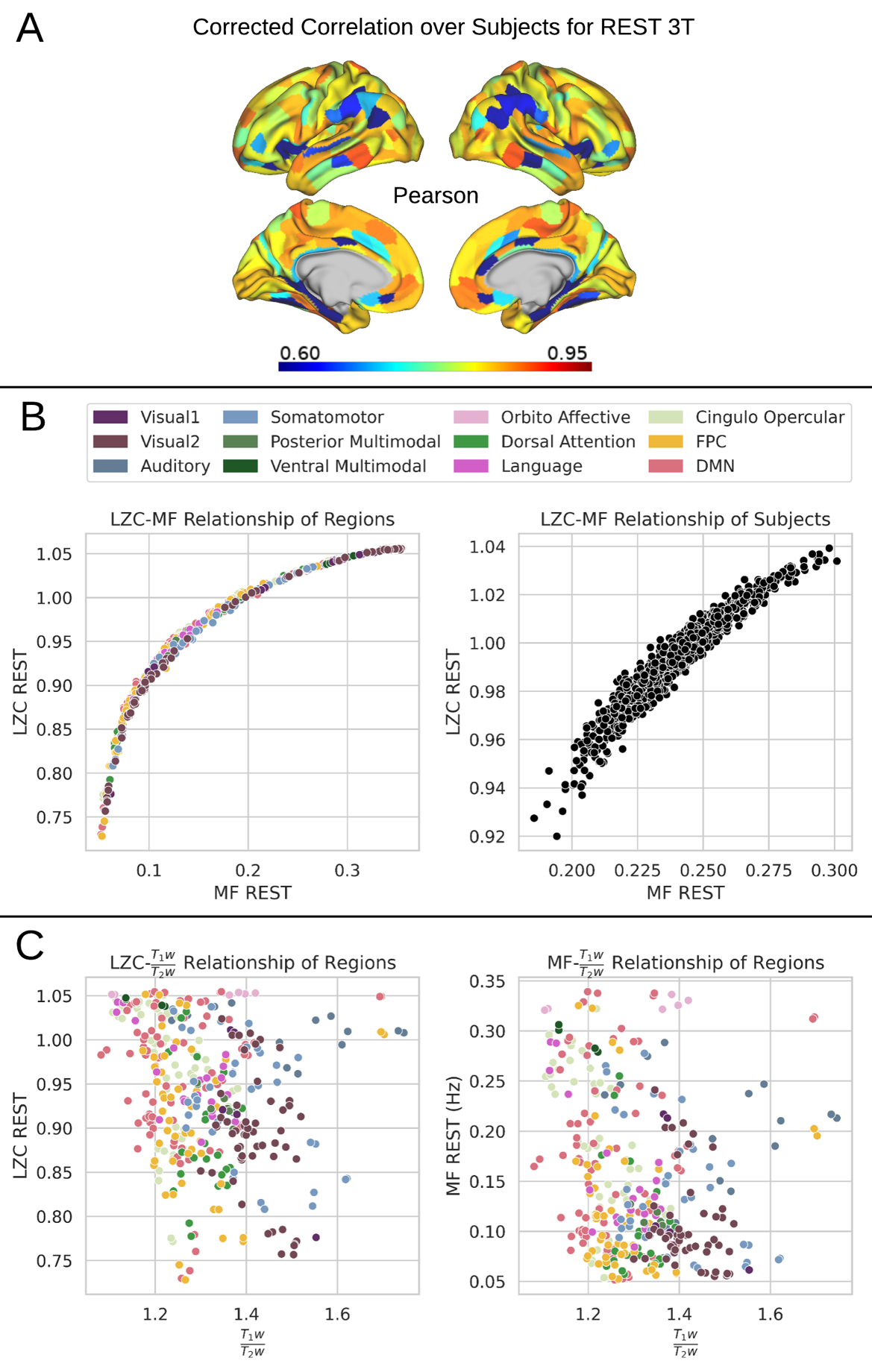

**Supp. Fig. 3. The relationship between LZC and MF during 3T resting state.** **(A)** Regional correlation between LZC and MF was computed by the Pearson method and corrected for multiple comparisons by the FDR method. **(B)** Regional scatter plots of LZC-MF relationship during 3T REST. LZC is plotted as a function of MF. Left shows scatter plot of regions averaged over subjects. Right shows subjects averaged over regions. **(C)** Scatter plots of LZC (left) and MF (right) as functions of structural ${T_{1}w}/{T_{2}w}$ $\frac{T_{1}w}{T_{2}w}$ values. Each point is a region averaged over subjects.

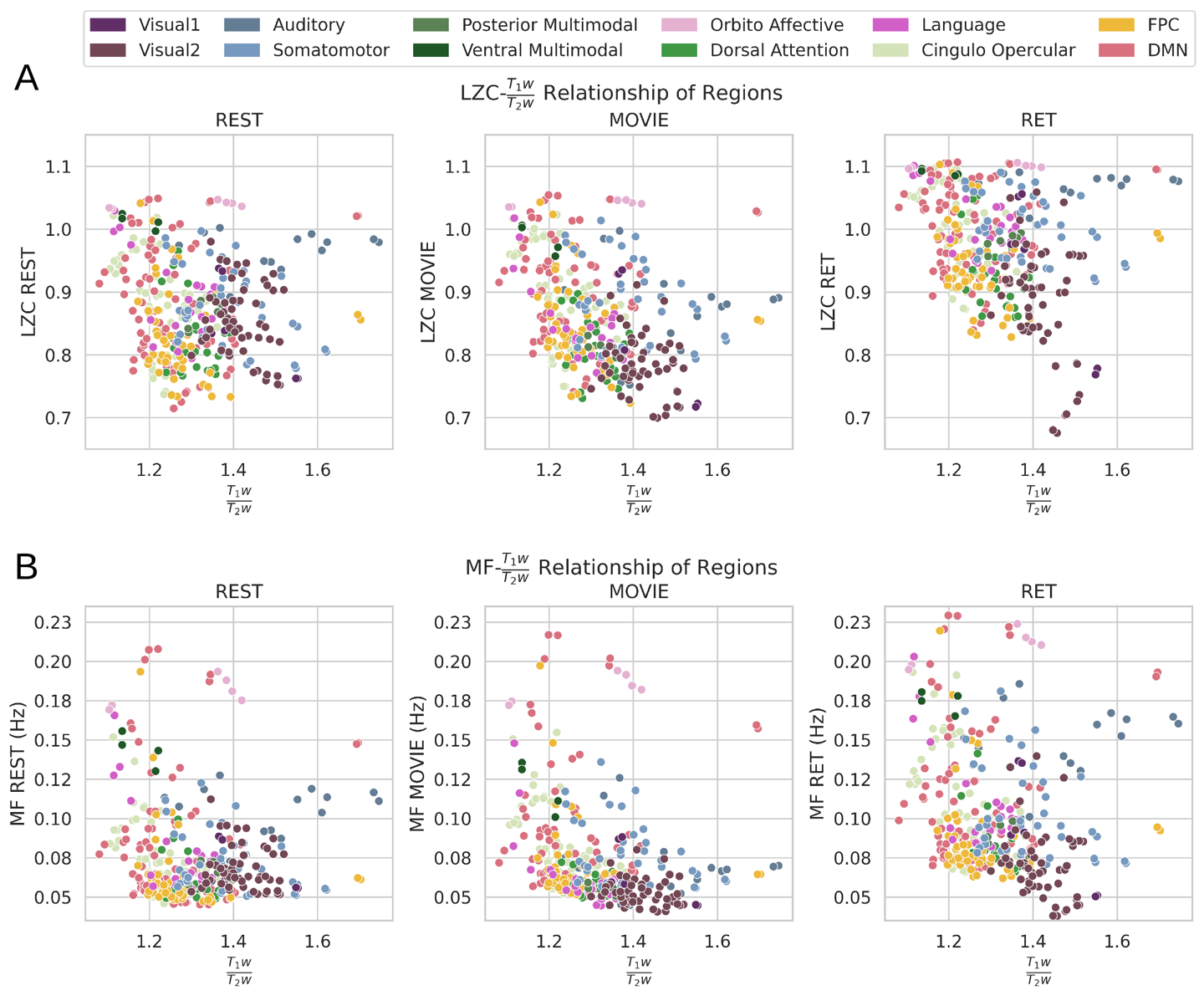

**Supp. Fig. 4. LZC/MF relationship with** ${T_{1}w}/{T_{2}w}$**.** The relationship between structural ${T_{1}w}/{T_{2}w}$ values and LZC/MF. The distribution of LZC (A) and MF (B) values were calculated for each region and plotted as functions of their corresponding ${T_{1}w}/{T_{2}w}$ values in all three conditions of REST (left), MOVIE (middle) and RET (right). No non-linear relationship can be observed.

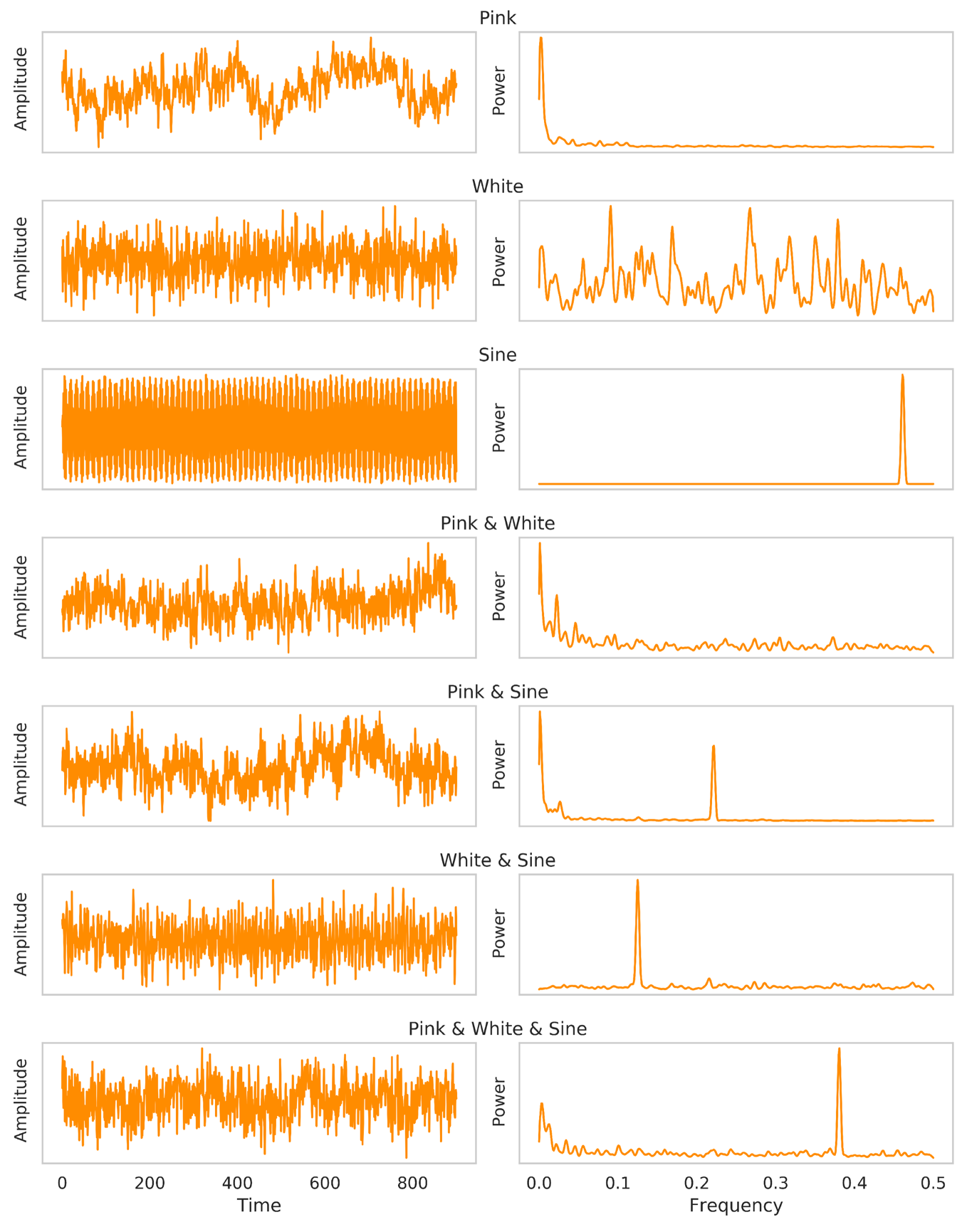

**Supp. Fig. 5. Sample simulated signals in each category.** Left plots are the sample signals in the time domain and the right ones are the same signals in the frequency domain (their power spectral distribution). For each category 5000 similar signals with different parameters were calculated and used in our LZC-MF relationship analysis.

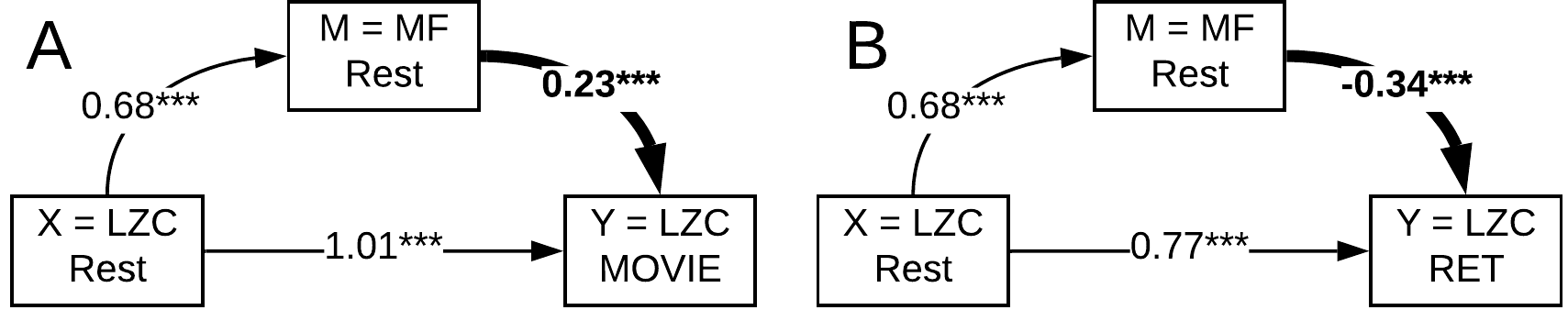

**Supp. Fig. 6.** Mediation model to investigate the role of MF-REST in the change in LZC from resting to task state. LZC-REST is used as the independent variable and the model investigates whether the effect of LZC-REST on LZC-MOVIE (A) or LZC-RET (B) as the dependent variable is mediated by MF-REST. The bootstrapped model showed significant partial mediation through MF-REST.
